# Matrix-trapped viruses can protect bacterial biofilms from invasion by colonizing cells

**DOI:** 10.1101/2020.10.28.358952

**Authors:** Matthew C. Bond, Lucia Vidakovic, Praveen K. Singh, Knut Drescher, Carey D. Nadell

## Abstract

Bacteria often live in the context of spatially restricted groups held together by a self-secreted, adhesive extracellular matrix. These groups, termed biofilms, are likely where many phage-bacteria encounters occur. A number of recent studies have documented that phages can be trapped in the outer matrix layers of biofilms, such that the bacteria inside are protected from exposure. It is not known, however, what might happen after this: are the trapped phages still viable on the biofilm exterior? If so, do they pose a threat to newly arriving cells that might otherwise colonize the existing biofilm? Here we set out to address these questions using a biofilm-producing strain of *Escherichia coli* and its lytic phage T7. Prior work has demonstrated that T7 phages are trapped in the outermost layers of curli polymers within the *E. coli* matrix. We show that these phages do remain viable and kill incoming colonizing cells so long as they are T7-susceptible. If colonizing cells arrive to the outside of a resident biofilm before phages do, they can still be killed by phage exposure if it occurs soon thereafter. However, if colonizing cells are present on the biofilm long enough before phage exposure, they gain phage protection via envelopment within curli-producing clusters of the resident biofilm cells. This work establishes that phages trapped in the outer matrix layers of a resident biofilm can be incidentally weaponized as a mode of protection from competition by newly arriving cells that might otherwise colonize the biofilm exterior.

## Introduction

Bacteria and their bacteriophage predators, or phages, are the most widespread biological entities on our planet. Each is found alongside the other in nearly every environment examined ^1–4^. Phages inject their genomes into the cytoplasm of their hosts, and in the case of obligate lytic phages, immediately begin co-opting host resources to replicate. Eventually, host cells are lysed to release a new cohort of phage virions. Predatory pressure from phage attack drives bacterial evolution, diversification, and ultimately the community structure of many microbiomes ^5–9^. The mechanistic, ecological, and evolutionary features of phage-bacteria interactions have a deep history of study, including many seminal theoretical and experimental papers that have characterized the population and evolutionary dynamics of phage-bacteria interactions ^8,10–12^. The traditional literature in this area mostly considers well-mixed culture conditions such as those in shaken liquids, which can reveal fundamental aspects of phage-bacteria interactions without spatial structure. However, in nature bacteria often reside in spatially constrained, surface-bound communities, or biofilms ^13–16^. The limited work that has focused on phage infection in biofilm environments has often found outcomes that differ substantially from those observed in mixed liquid environments ^17–24^.

A defining feature of biofilm populations is the presence of a self-secreted adhesive polymer matrix that modulates cell-cell and cell-surface interactions, in addition to influencing collective biofilm architecture ^13,15,25–28^. Several recent papers have demonstrated that the biofilm matrix can be central to phage-host coevolution. *Pseudomonas fluorescens* and *Escherichia coli* rapidly evolve mucoid colony phenotypes – which reflect increased and/or altered matrix secretion – when they are under phage attack ^29,30^. Curli fibers, a proteinaceous component of the *E. coli* matrix, can block biofilm-dwelling cells from T7 and T5 phage exposure ^31^. In this case phages remain suspended in the curli mesh without infecting biofilm cells unless the integrity of the curli layer is compromised. Recent work by Darch et al.^32^, Diaz-Pascual et al.^33^, and Dunsing et al.^34^, respectively, suggest a similar pattern of matrix-dependent phage protection in *Pseudomonas aeruginosa*, *Vibrio cholerae*, and *Pantoea stewartii*. In those works which documented phage trapping in the extracellular matrix, it was unclear whether matrix-entangled phages are neutralized or remain infectious.

Inspired by the findings above, we investigated the consequences of phage entrapment in the matrix for biofilm population assembly. If they remain infectious, these phages could pose a threat to new cells that attempt to colonize the biofilm surface. Here we set out to explore this possibility by studying how matrix-embedded phages influence the invasion of bacteria into pre-existing biofilm populations, and whether biofilm-invading cells can integrate into the existing matrix and gain phage protection. We find that matrix-trapped phages do indeed remain viable and can dramatically shift the ability of newly arriving cells to colonize existing biofilms, and that this effect is dependent on the relative timing of the arrival of phages and colonizing cells to resident biofilms.

## Results and Discussion

### Influence of matrix-trapped phages on invading planktonic cells

*E. coli* produces a variety of matrix components at different times during biofilm formation, including flagellar filaments, curli amyloid fibers, and polysaccharides including colanic acid and cellulose ^31,35–41^. Curli fibers are the single matrix component known to be essential for T7 phage protection in *E. coli* biofilms ^31^ and are assembled by extracellular polymerization of CsgA monomers on an outer membrane baseplate comprised of CsgB ^37,42^. Curli proteins secreted by *E. coli* localize primarily in the upper half of the biofilm-dwelling population ^31,40,41^, and introduced phages become trapped in the outer part of this curli matrix layer (Figure 1A). These phages are blocked from diffusing into the biofilm interior, but we hypothesized that trapped phage particles may remain capable of infecting cells that reach them from the liquid environment that surrounds the biofilm. To assess this question, we first grew biofilms of *E. coli* AR3110 for 62 hours (Figure 1B). The resident cells harbored a chromosomal construct for constitutive expression of the far-red fluorescent protein mKate2^43^ to make them visible with fluorescence time-lapse microscopy. The biofilms were then exposed to a 1 h pulse of media containing lytic T7 phages at 2×10^9^ PFU/mL, such that phages could accumulate in the outer matrix curli layer (Figure 1C). The phages contained a *sfGFP*^44^ expression construct engineered into their genome, allowing us to visualize any cells that became phage-infected^31^. For a control comparison, we performed the same procedure but exposed the resident biofilms to sterile media containing no phages. Following phage exposure or control treatment, we performed a population invasion step in which high-density planktonic cultures of phage-susceptible wild type *E. coli* AR3110 (5 × 10^9^ CFU/mL) were added to the chambers for 3 hours to colonize the resident biofilms (Figure 1D). The experimental procedure for these experiments is summarized in Figure S1. Colonizing cells were isogenic to resident wild type cells, except they constitutively expressed mKO-κ^45^ so that they could be distinguished from resident cells during imaging.

**Figure 1:**
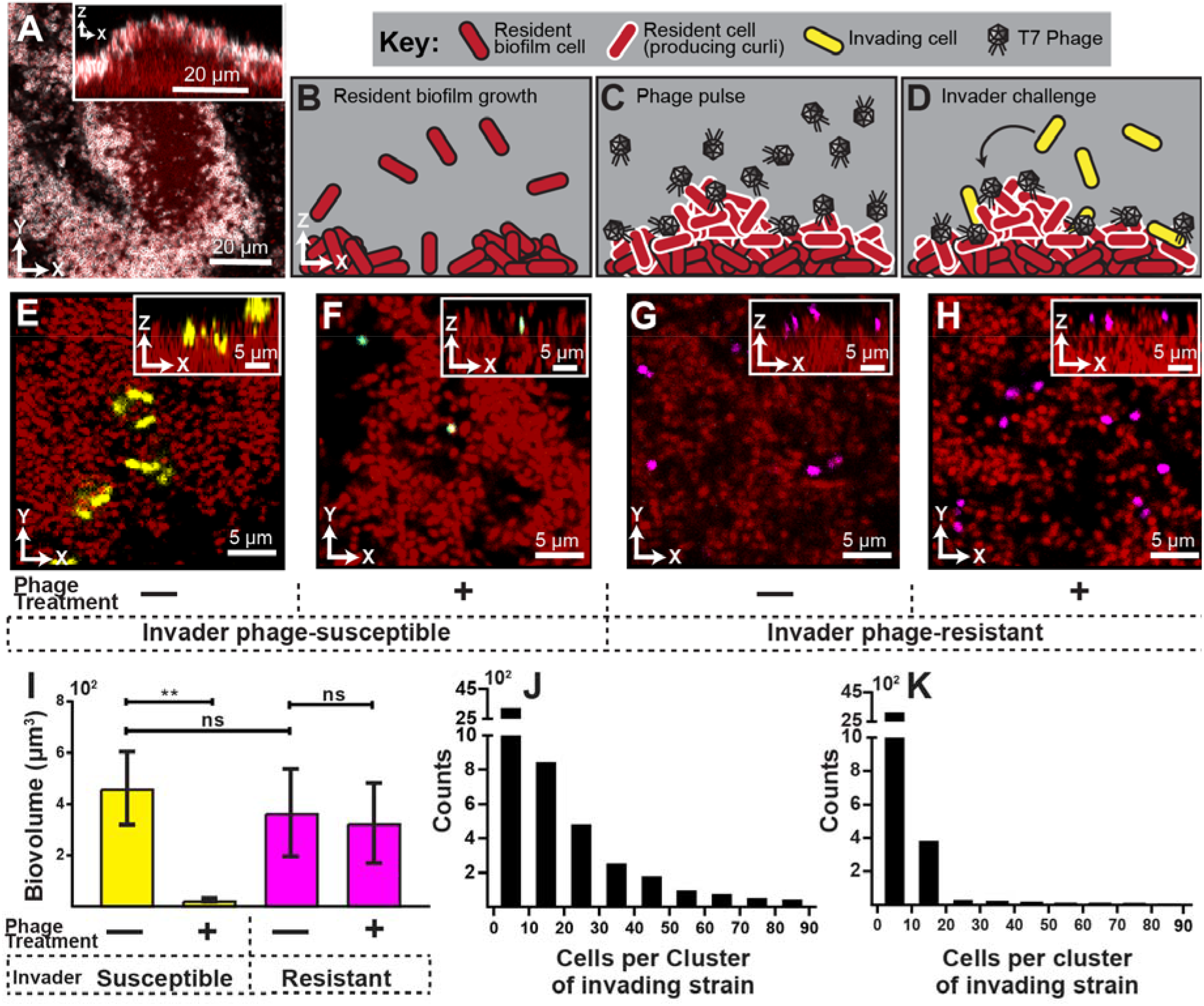
Visualization and quantification of biofilm invasion with or without phage exposure. (A) Visualization of *E. coli* biofilm (red) with stained curli matrix (white), including one optical section (main image) and z-projection (inset). (B-D) Illustration of invasion assay procedure. (B) Schematic of resident biofilm growth (red cells). (C) Inlet tubing is then swapped for new tubing & syringe for 1 h, containing a highly concentrated phage suspension. (D) Resident biofilms were then challenged with isogenic *E. coli* expressing a different fluorescent protein (yellow). Inlet tubing was then again swapped to a new tubing & syringe for 1 h, containing high density *E. coli* culture. Biofilms were imaged for 10 hours following this step. (E) Invading cells (yellow) successfully attach to the resident biofilm (red) in the absence of phages. (F) Invading cells fail to integrate when biofilms are pretreated with phages, which become trapped in the biofilm matrix. Phage-encoded fluorescent infection reporter (blue) indicates invading cells that have become phage-infected. (G) A phage-resistant mutant (Δ*trxA*, magenta) invades with equivalent frequency in control and phage pre-treatment conditions. (I) Quantification of image data shown in E-H; average invading biovolume per field of view (150 μm x 150 μm x 15 μm; L, W, H). Error bars represent standard error of the mean. Pairwise comparisons were performed with Mann-Whitney Signed Ranks tests with Bonferroni correction (*n*=6; ** denotes *p*<0.005). (J-K) Invading cell cluster size distributions for phage-susceptible cells invading biofilms without (J) or with (K) phage pre-treatment.

We found that in the absence of phage exposure, resident biofilms could be colonized by planktonic cells, albeit not at high efficiency (Figure 1E, I). The colonizing cells were restricted to the outer surface of the resident biofilms and could not enter the biofilm interior, similar to what we have seen previously for *V. cholerae*^46^. However, when resident biofilms were pre-exposed to phages, colonization of phage-susceptible cells of the resident biofilms was almost completely eliminated (Figure 1F, I). Further, most invading cells that were detected on phage-exposed biofilms were fluorescent in the sfGFP channel, indicating that they had been phage-infected but not yet lysed. Our interpretation of this result is that susceptible invading cells encounter phages in the curli matrix, become infected, and lyse or fail to divide further. Another (though not mutually exclusive) possibility is that phages, by occupying potential sites of attachment, block the physical interaction of invading cells with the biofilm outer surface. We tested this possibility by repeating the experiment above with invading cultures of an *E. coli* mutant harboring a clean deletion of *trxA*, which confers resistance to T7 phage infection ^47^. The Δ*trxA* deletion mutant was found to invade resident biofilms at equal rates whether or not the resident was pre-exposed to T7 phages. This result eliminates the possibility that phages block attachment sites for invading cells (Figure 1G, H, I).

### Invading cells gain phage protection, after a delay

Once we determined that matrix-embedded phages could infect recently attached susceptible cells, we asked whether invading cells fare better if they arrive prior to the phage pulse. To explore this question, we again grew resident biofilms of *E. coli* AR3110 for 62 h, followed by a planktonic population invasion step as described above. Chambers were separated into two groups: In the first group, phages were pulsed into the chamber immediately after colonization by the invading cell population. In the second group, biofilms were incubated post-invasion for 10 h, then subjected to a phage pulse. This experimental procedure is summarized in Figure S2. Images were taken of each chamber approximately 10 h after the phage pulse, allowing for multiple infection and lysis cycles to occur. When phage pulses occurred immediately after biofilms were colonized by an invading strain, the invading cells were mostly killed by T7 phage exposure (Figure 2A, B). Interestingly, however, invading cells were not infected by phage pulses that arrived 10 h after the colonizing strain attached to the resident biofilm outer periphery (Figure 2A, C).

**Figure 2:**
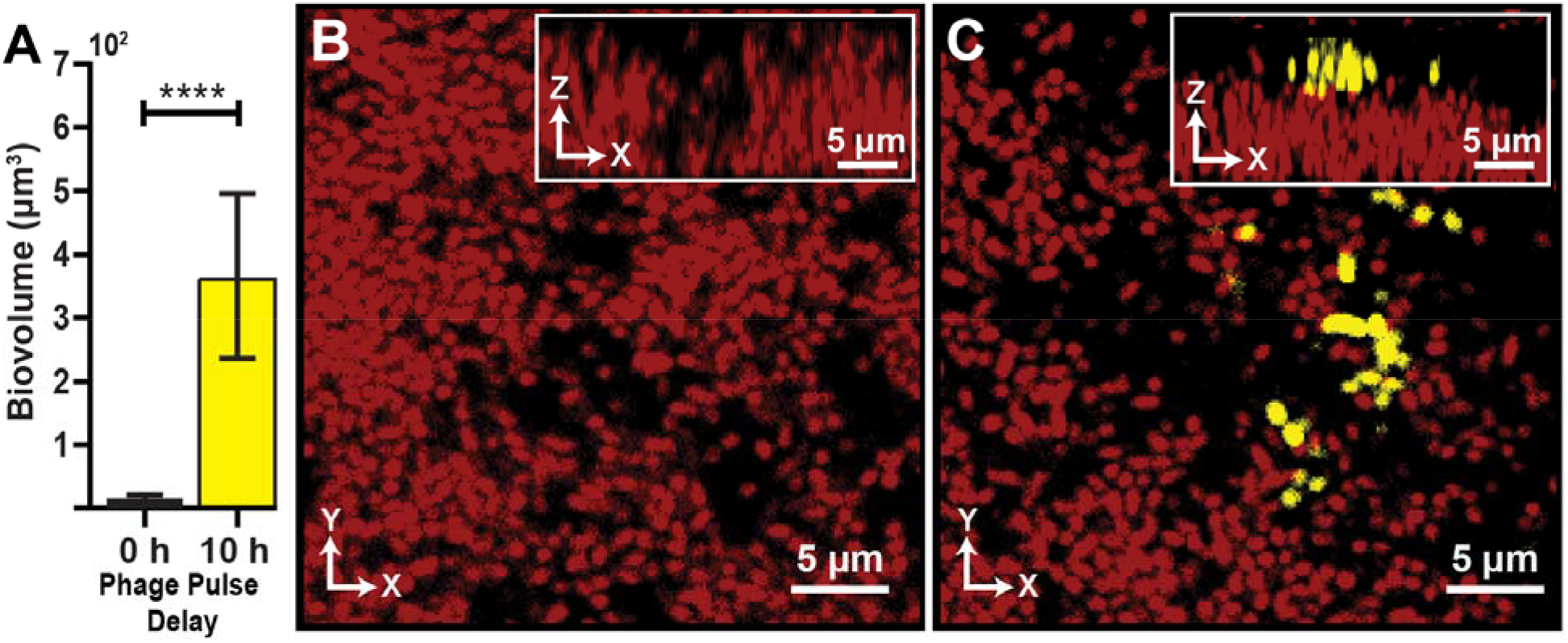
Visualization and quantification of colonization success with phage exposure post-colonization. (A) Average invading biovolume per field of view (150 μm x 150 μm x 15 μm; L, W, H). Error bars represent standard error of the mean. Pairwise comparisons were performed with a Mann-Whitney Signed Ranks test (*n*=7; **** denotes *p*<0.00005). (B) Invading cells (yellow, though absent in B) are killed when phages are introduced immediate after their arrival. Resident biofilm cells Are shown in red (C) Invading cells are not killed when phages are introduced 10 h after their arrival.

### Invading cells indirectly co-opt matrix of resident biofilm

Having observed that invading cells gain phage protection, but only after a delay between attaching to a resident biofilm and phage exposure, we next investigated how invading cells gain phage protection over this delay period. Given that phage protection is dependent on being embedded in curli polymers of the biofilm matrix ^31^, it is possible that colonizing cells produce their own curli after an initial transition period following attachment to the resident biofilm. Alternatively, invading cells could co-opt matrix produced by the resident population. We tested these possibilities using a combination of experiments in which either the invading strain or the resident strain produced a 6xHis-labeled variant of CsgA, allowing us to localize and quantify curli production as a function of time and space inside biofilms via immunostaining. We cultivated resident biofilms and performed the invasion assay as above (summarized in Figure S2), but in this case we included an anti-His, AlexaFluor-647-conjugated antibody in the influent medium such that any curli produced in the biofilm became fluorescent and detectable by confocal microscopy (Figure 3A).

**Figure 3:**
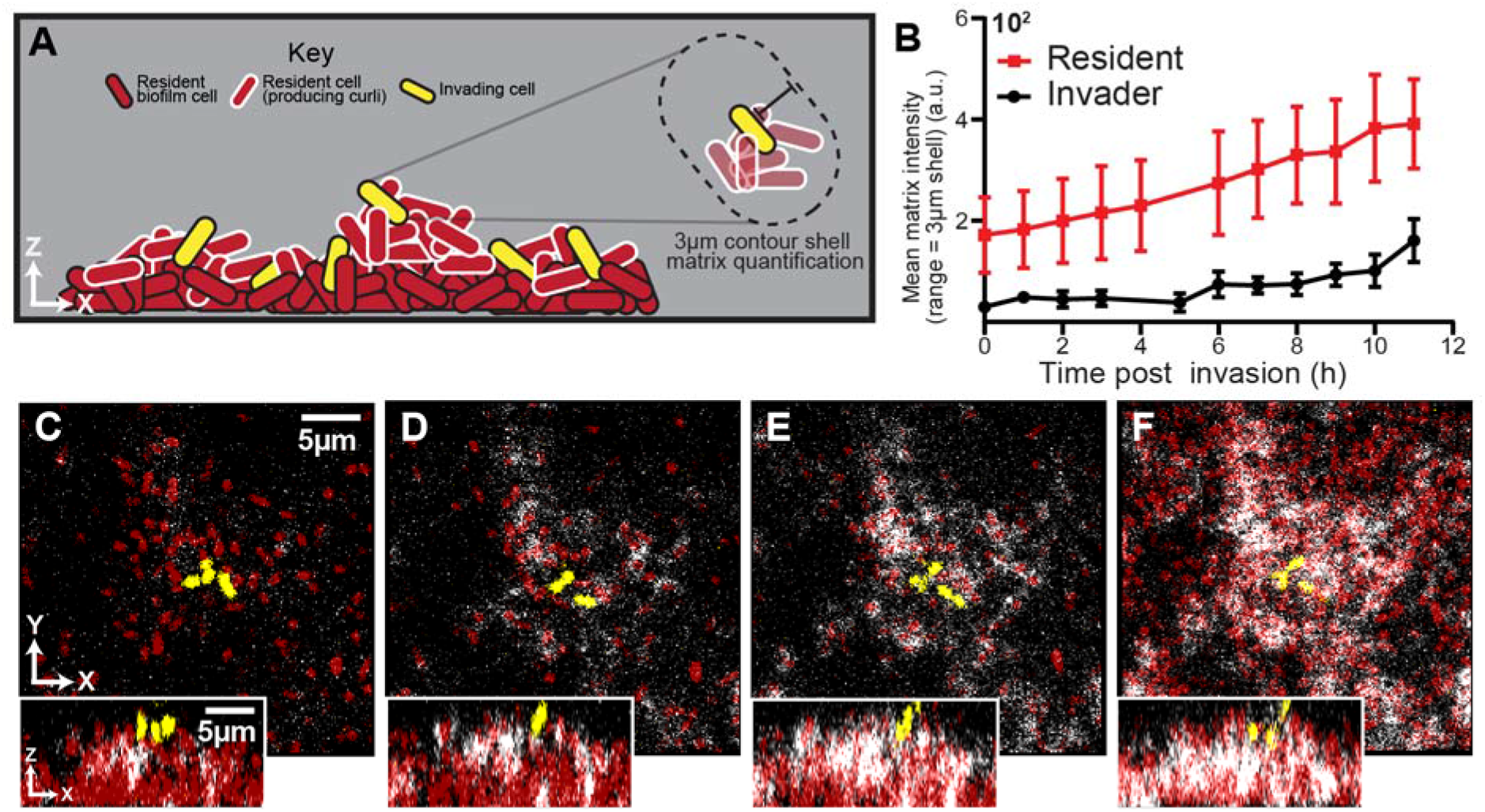
Spatial and temporal dynamics of curli fiber localization around resident and invading cells. (A) Illustration of matrix quantification method. Fluorescence intensity of labeled curli fibers is determined in a 3 μm shell surrounding individual segmented cell volumes. (B) Quantification of matrix localization surrounding the resident and invading populations during an 11 h time course after invading cells arrive (*n*=4, errors bars denote SE). (C-F) Representative images displaying labeled curli matrix (white), invading cells (yellow) and the resident, matrix-producing cells (red). Images were acquired at 1 h, 5 h, 7 h and 10 h following the arrival of invading cells.

When invading cells harbored the His-labelled variant of *csgA*, we detected negligible anti-His fluorescence in the 10 hours following invasion of the biofilm exterior (SI Figure S3). By contrast, when cells in the resident biofilm harbored the His-labelled variant of *csgA*, abundant anti-His staining was detectable in the upper surface of the resident biofilm (Figure 3 B, C), including the surroundings of the colonizing cells ^31,40,41^. These two observations indicate that invading cells do not gain phage protection in the 10 hours following colonization because they produce their own curli fibers, but rather because they coopt the curli fibers being produced by cells in the resident biofilm (Figure 3B). To corroborate this interpretation we used image analysis to quantify curli accumulation in the immediate neighborhood of resident and colonizing cell biomass following invasion; this analysis showed a steadily increasing amount of curli in the immediate neighborhood of colonizing cells over the course of 10 hours following their arrival to the outer surface of the resident biofilm. At the 10-hour mark, invading cells had the same amount of curli around them as the resident strain did at the beginning of the invasion experiment.

We next asked if the invading strain directly or indirectly co-opts curli matrix material from the resident biofilm. Invading cells producing CsgB baseplates could potentially collect freely diffusing CsgA monomers being released by the resident biofilm population, accumulating curli matrix material and thus phage protection. This might not necessarily be the case, as our experiments above showed that the invading cells were not producing CsgA, whose corresponding gene is in the same operon as *csgB*; one would typically not expect the production of one without the other. Alternatively, without directly sequestering resident-produced CsgA to their cell surface, invading cells could become enveloped within curli-producing clusters of the resident biofilm population with enough curli in their surroundings to block phage diffusion. This would be an indirect way of exploiting the phage protection of the resident biofilm’s curli layer.

To differentiate between these two possibilities, we repeated the post-invasion phage pulse experiments using a Δ*csgB* mutant that cannot produce its own CsgB base plate, and which therefore cannot nucleate CsgA polymerization on its outer surface. Here, as above (see Figure 2), few cells survived if phages arrived immediately after invading cells colonized the resident biofilm surface (Supplemental Figure S4). However, despite lacking CsgB curli baseplates, invaders were still protected after a 10-hour delay prior to phage exposure in the system, and their accumulated biovolume was not statistically different from that of a wild type invading strain (Supplemental Figure S4). This outcome suggests that invading cells do not have to be able to directly polymerize CsgA on their exterior to gain phage protection over the 10 hours after attaching to the biofilm surface. We interpret this result to indicate that the invading cells *indirectly* coopt curli produced by the resident biofilm by becoming sufficiently enveloped amidst clusters of resident cells that their exposure to incoming phages is greatly reduced or eliminated.

## Conclusions

We have demonstrated that matrix-embedded phages can remain infectious on the biofilm surface and kill newly arriving bacteria. In this sense, biofilm-dwelling microbes, by trapping phages on their exterior, can weaponize them against incoming phage-susceptible cells. On the other hand, if invading *E. coli* attach to a resident biofilm and have sufficient time to become entangled in the curli matrix, they too gain protection from subsequent phage exposure. This protection is obtained by indirect exploitation of the resident biofilm’s curli matrix: invading cells did not significantly use resident-produced curli monomers to polymerize curli on their own exterior, but rather became sufficiently enveloped in curli-producing groups of resident biofilm cells that they were no longer exposed to an incoming phage attack.

Our observations bear an interesting analogy to those results of Barr et al.^48^, who found that phages trapped in host mucosal linings can kill incoming bacteria. We speculate on the basis of our results here that phage entrapment and their blocking effect against bacterial colonization is important not just in host associated mucosal environments but even more broadly to many biofilm contexts in which phage-trapping matrix material could potentially influence the pattern of community succession.

As a proof of principle, we studied here the effect of matrix-embedded phages on the colonization ability of cells isogenic to those in the resident biofilm, showing that biofilm colonization was effectively eliminated by the presence of phages on the biofilm surface. We speculate that it is unlikely that phage-trapping on biofilm surfaces is unique to *E. coli*. Recent reports from other groups have suggested matrix-dependent protection against and potentially sequestration of incoming phages, but it remains to be tested whether phages remain active against other incoming cells in these cases^32–34^. The centrally important question prompted by our results here is whether biofilm consortia trap phages of many different strain and species specificities, each of which has the potential to kill off invading competitors of its target host range. In this sense the biofilm matrix, among its many other functions, may protect the cells within against spatial competition with many other species *via* phage entrapment.

## Materials and Methods

### Strains

*E. coli* strains used in these experiments were all derived from *E. coli* AR3110 and are listed in Supplemental Table 1. Each strain used for imaging contained a codon-optimized fluorescent protein construct (*mKO-κ*, *mKate2* or *mTFP1*) under the control of a constitutive P_taq_ promoter. Fluorescent protein constructs were inserted using the traditional lambda-red recombineering method by amplifying constructs with primer overhangs corresponding to the *attB* site (26). iProof High-Fidelity DNA Polymerase (Bio-Rad, Hercules, CA, USA) was used to amplify insertion sequences for fluorescent protein expression constructs. The *E. coli* Δ*trxA* deletion strain was also constructed using lambda-red recombineering. The deletion construct was made by amplifying a kanamycin resistance marker flanked by FRT sites. Primer overhangs added upstream and downstream regions flanking start and stop codons of the *trxA* locus for replacement of the full reading frame with the Kan cassette. FRT recombinase was subsequently expressed *in trans* to remove the kanamycin selection marker.

### Phage propagation and titer

T7 lytic phage was propagated and lysates collected in a manner adapted from Bonilla et. al ^50^. Briefly, AR3110 wild type *E. coli* cultures were grown overnight in 5 mL lysogeny broth at 37 °C at 250 rpm in a New Brunswick orbital shaking incubator. The host strain was then back diluted 1:20 into 100 mL lysogeny broth and allowed to grow to mid exponential phase (0.3-0.5 OD^600^). At this time, T7 phage were spiked in from a frozen stock and MgSO_4_ was added to a final concentration of 5 mM. The culture was placed back into the incubator for 3 to 4 hours, until the culture clarified. The entire volume was then vacuum filtered (0.22 μm filter Millipore Sigma). Phage titer was determined by traditional plaque assay^51^. Briefly, host *E. coli* was grown overnight and sub-cultured as described above to achieve mid exponential phase (0.3-0.5 OD^600^). Phage preparation was serially diluted by passing 10 μL into 990 μL for 100-fold dilutions. Top agar (0.5% agar, lysogeny broth) was melted and aliquoted into 3 mL volumes. Subsequently, 50 μL of a dilution was added to each sample along with MgSO_4_ (5 mM). Molten top agar was then poured evenly onto lysogeny broth plates and placed at 37 °C for 3 hours. Plates were removed and plaques were counted in order to back calculate plaque forming units (PFU) per milliliter.

### Biofilm phage pretreatment invasion assay

We measured attachment and growth of exogenously added planktonic cells to curli protected biofilms with and without the prior addition of T7 phage. *E. coli* AR3110 expressing mKate2 were cultured in 5 mL lysogeny broth overnight at 37 °C at 25 rpm in a New Brunswick orbital shaking incubator. Cultures were then pelleted and washed twice with 0.9% NaCl and standardized to 0.2 OD^600^ prior to inoculation into microfluidic devices. The inoculum was incubated for 1 h at room temperature (approximately 22°C). Media syringes (1 mL BD plastic) were prepared by loading 1mL of 1% tryptone broth (W/V) and attaching a 25-gauge needle. Tubing (#30 Cole palmer PTFE ID 0.3mm) was then carefully attached to the needle and syringes were subsequently placed in Harvard Apparatus syringe pumps. After affixing inlet and outlet tubing to the microfluidic devices, a 40 second pulse at 40 μL/min was conducted to remove unattached cells, before standard flow regime (0.1 μL/min) was started. Biofilms were grown at room temperature for 62 hours at which time, tubing was swapped from clean media to either purified phages (2×10^8^ PFU/mL) or clean media control. Flow was continued at 0.1 μL/min for 1 h. After phage pretreatment, tubing was again removed and switched to syringes containing invading cells expressing mKO-κ for three hours at 0.1 μL/min. Invading cells were prepared prior to this step. Cells were grown overnight as previously described before in lysogeny broth. Invading cells were then sub-cultured 1:20 into 100 mL of 1% tryptone broth for 3 hours at 37°C at 250 rpm. Cells were then pelleted, concentrated, and standardized to 6.0 OD^600.^ Following the conclusion of the invasion, microfluidic chambers were allotted 10 hours of incubation at room temperature under standard flow conditions for growth and phage infection to occur. Three to five image fields (150 μm x 150 μm x 25 μm; L x W x H, slice interval 0.5 μm) were then acquired on a Zeiss 880 LSCM and image analysis performed using BiofilmQ software tool ^52^.

### Biofilm phage posttreatment invasion assay

Attachment and growth of invading *E. coli* were measured under two different regimes. Invasions of resident biofilms were conducted prior to phage application in this assay; however, timing of the phage application was varied. Resident biofilms were cultured in the same manner described above in the pretreatment invasion assay. At 62 hours of growth, clean media tubing was exchanged for invading cells (prepared in an identical manner as above) and allowed to flow for three hours at 0.1 μL/min. Following the conclusion of the invasion, chambers were separated into two groups corresponding to phage treatment regime. Half of the chambers were exposed to phage treatment (2×10^8^ PFU/mL at 0.1 μL/min for 1 h) immediately following the invasion, while the other half was incubated for 10 hours at room temperature prior to the phage treatment. In both regimes, images were acquired in the manner described above. Imaging took place 10 hours following their respective phage treatments.

### Curli matrix localization assay

Curli matrix monomers, CsgA, were labeled with a 6X-His tag as previously published^31^. Curli fibers were detected via direct fluorescent immunostaining with an α-6X-His antibody (Invitrogen) conjugated to a fluorescent dye, Alexafluor647. Antibody was added to clean media at 0.1mgL/mL and flowed continuously throughout the course of the experiment. Biofilms were grown as described above, using *E. coli* expressing labeled curli.

### Confocal microscopy and image analysis

All imaging was performed using a Zeiss 880 line-scanning confocal microscope with either a 10x/0.4NA or a 40x/1.2NA water objective to minimize axial aberration effects. The sfGFP fluorescent protein was excited using a 488 nm laser line, the mKO-κ fluorescent protein was excited using a 543 nm laser line, the mKate2 fluorescent protein was excited using a 594 nm laser line, and Alexafluor647 was excited using a 633 nm laser line. All image stacks were trimmed if necessary (e.g., if area outside of the microfluidic devices had been acquired in addition to the biofilm itself) using the native Zeiss Zen Blue software. All subsequent quantification was performed using the BiofilmQ image analysis tool framework^52^.

## Acknowledgments

CDN conceived and supervised the project. CDN and MCB designed experiments. MCB performed experiments and image analysis. PKS, LV, and KD provided critical reagents. CDN, MCB, and KD finalized figures. CDN and MCB wrote the paper with input from all authors.

## Supplemental Information

**Supplemental Table S1.**
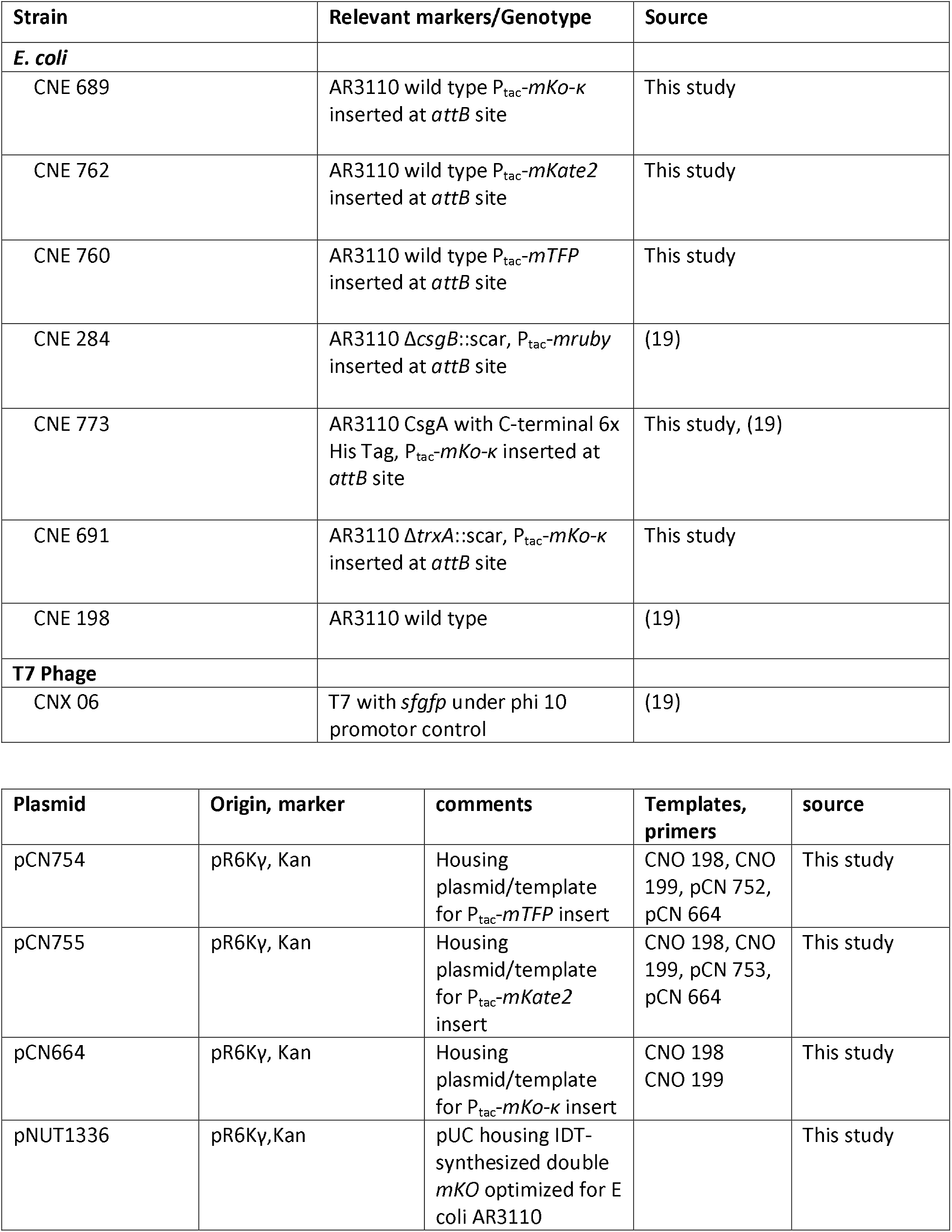
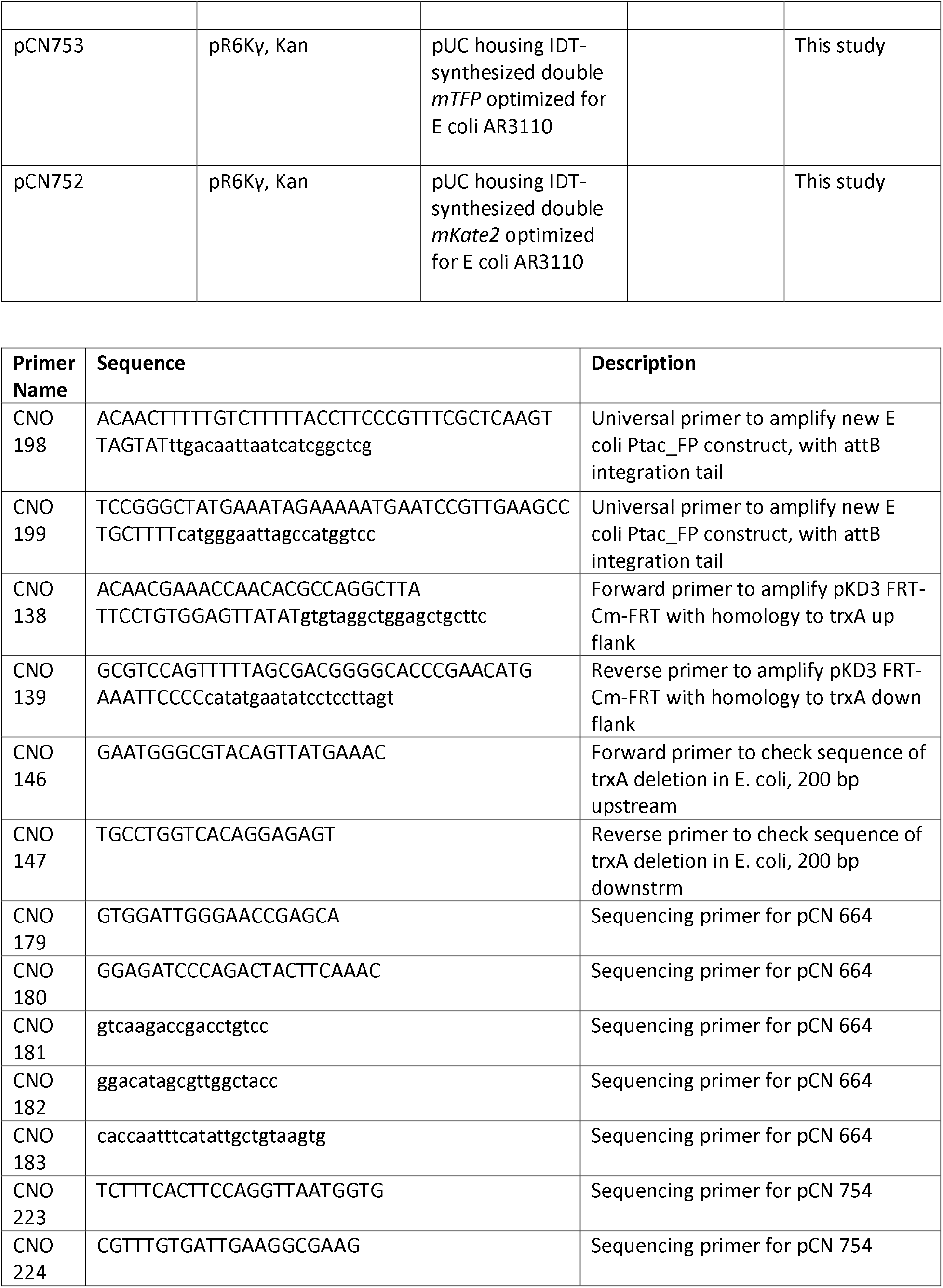
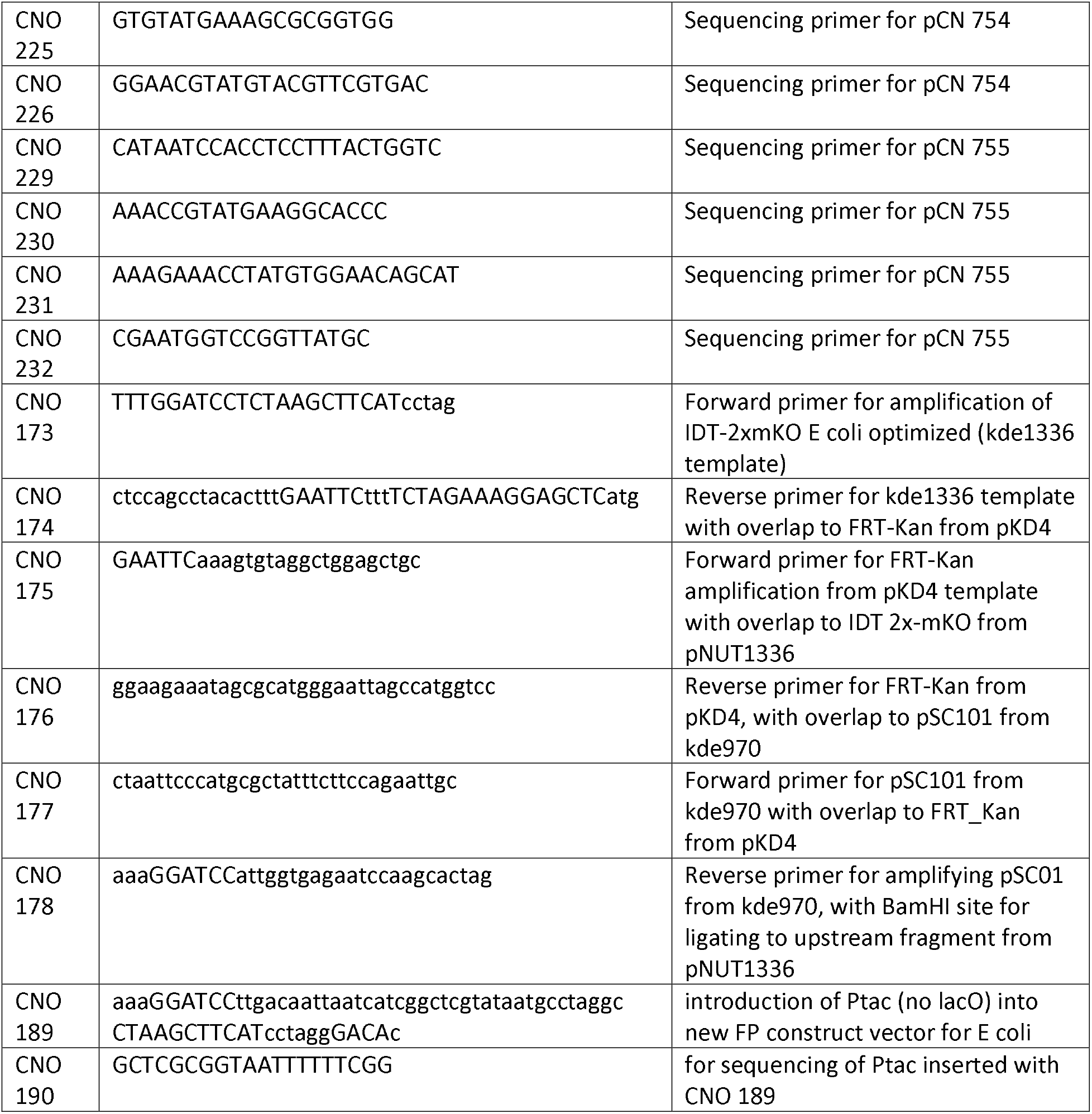
Strains, plasmids, and oligos used in this study.

## Supplemental Figures

**Figure S1:**
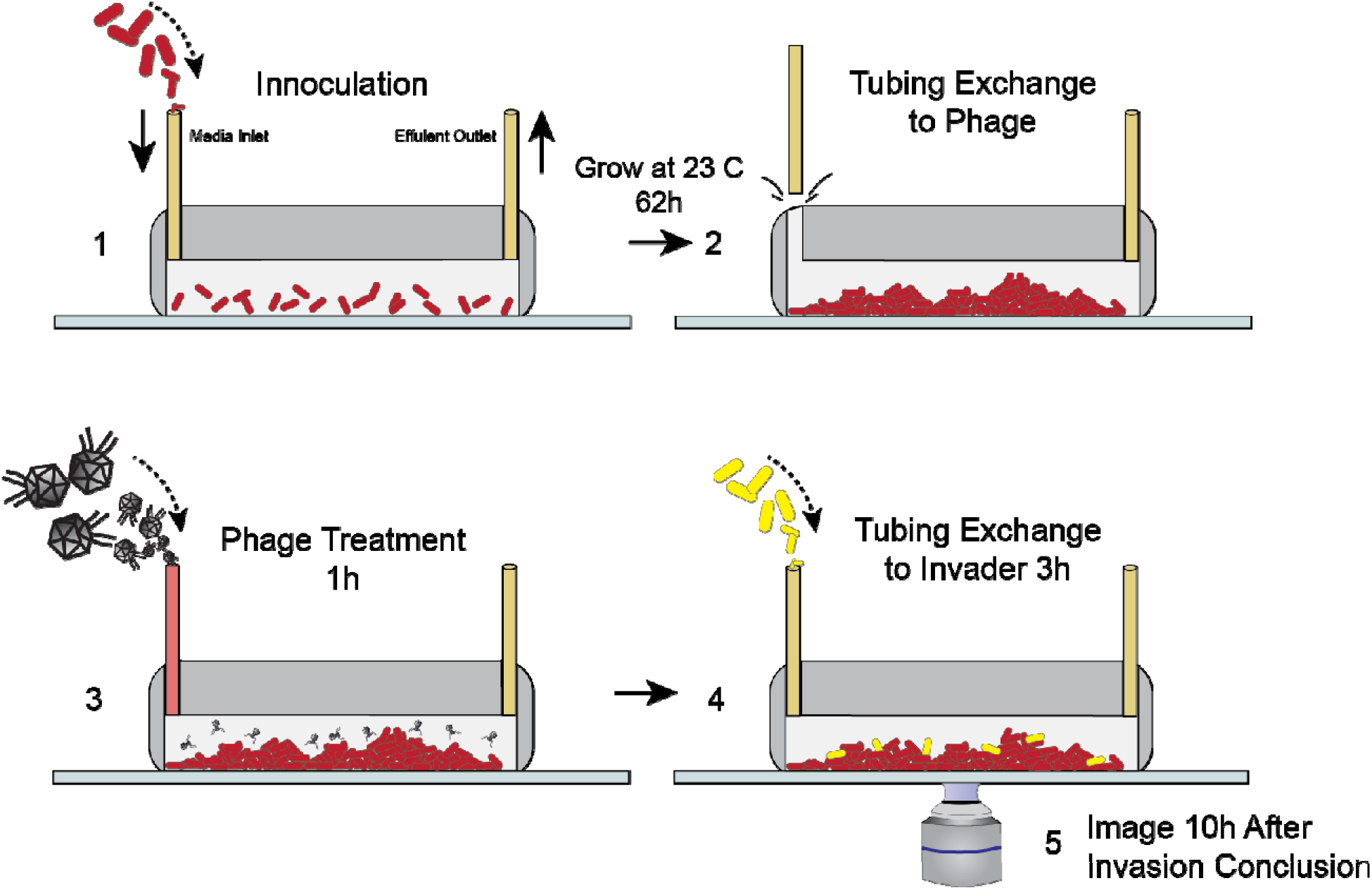
Method for Phage Pretreatment Experiments. (A) *E. coli* resident strain was inoculated into microfluidic devices at OD^600^ = 0.2. Cells were allotted an hour for surface attachment before a media flush was performed (40 seconds). Biofilms were then grown at room temperature for approximately 62 hours. (B) Inlet **t**ubing was carefully removed from device and replaced with tubing containing phages. (C) Phages (2×10^9^ PFU/mL) were flowed into the chamber for 1 hour (0.1 μL/min for 1h). (D) Following phage pretreatment, isogenic *E. coli* invaders (OD^600^ 6.0) were flowed into the chamber, again through a tubing swap. (E) Tubing was finally returned to sterile media and the device was allowed to incubate for 10 h prior to imaging.

**Figure S2:**
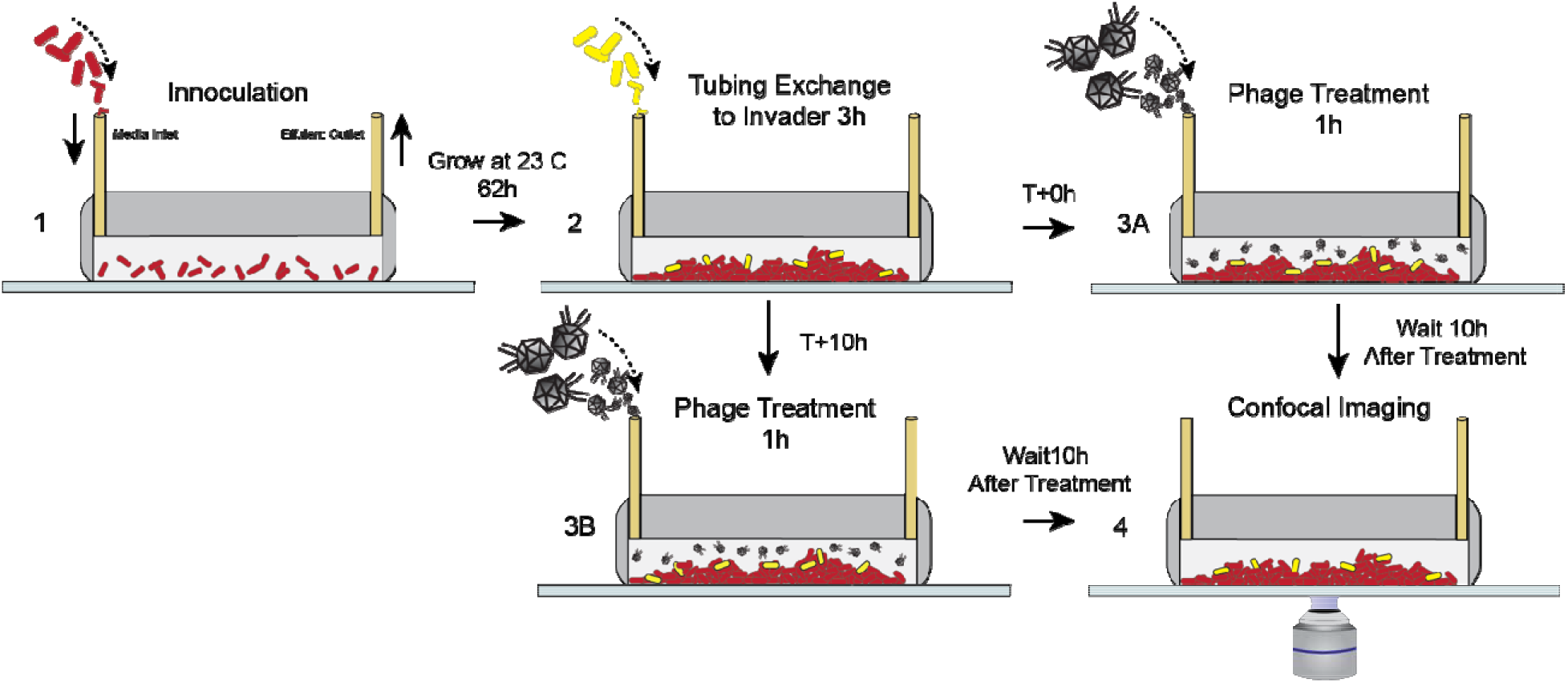
Method for Phage Posttreatment Experiments. (A) *E. coli* resident strain was inoculated into microfluidic devices at OD^600^ 0.2. Cells were allotted an hour for surface attachment before a media flush was performed (40 seconds). Biofilms were then grown at room temperature for approximately 62 hours. (B) Invading cells were then flowed into all chambers (OD^600^ 6.0) for three hours. After the invasion, individual chambers were separated into two groups: (C,D) Chambers in group **C** received phage treatment (2×10^9^ PFU/mL, 0.1 μL/min for 1h) immediately following the invasion. These chambers were imaged ten hours after the conclusion of the phage treatment (**S1E**). Chambers in group **D** were incubated at room temperature for ten hours before receiving phage treatment. Ten hours following the completion of the phage treatment, chambers in group **D** were imaged.

**Figure S3:**
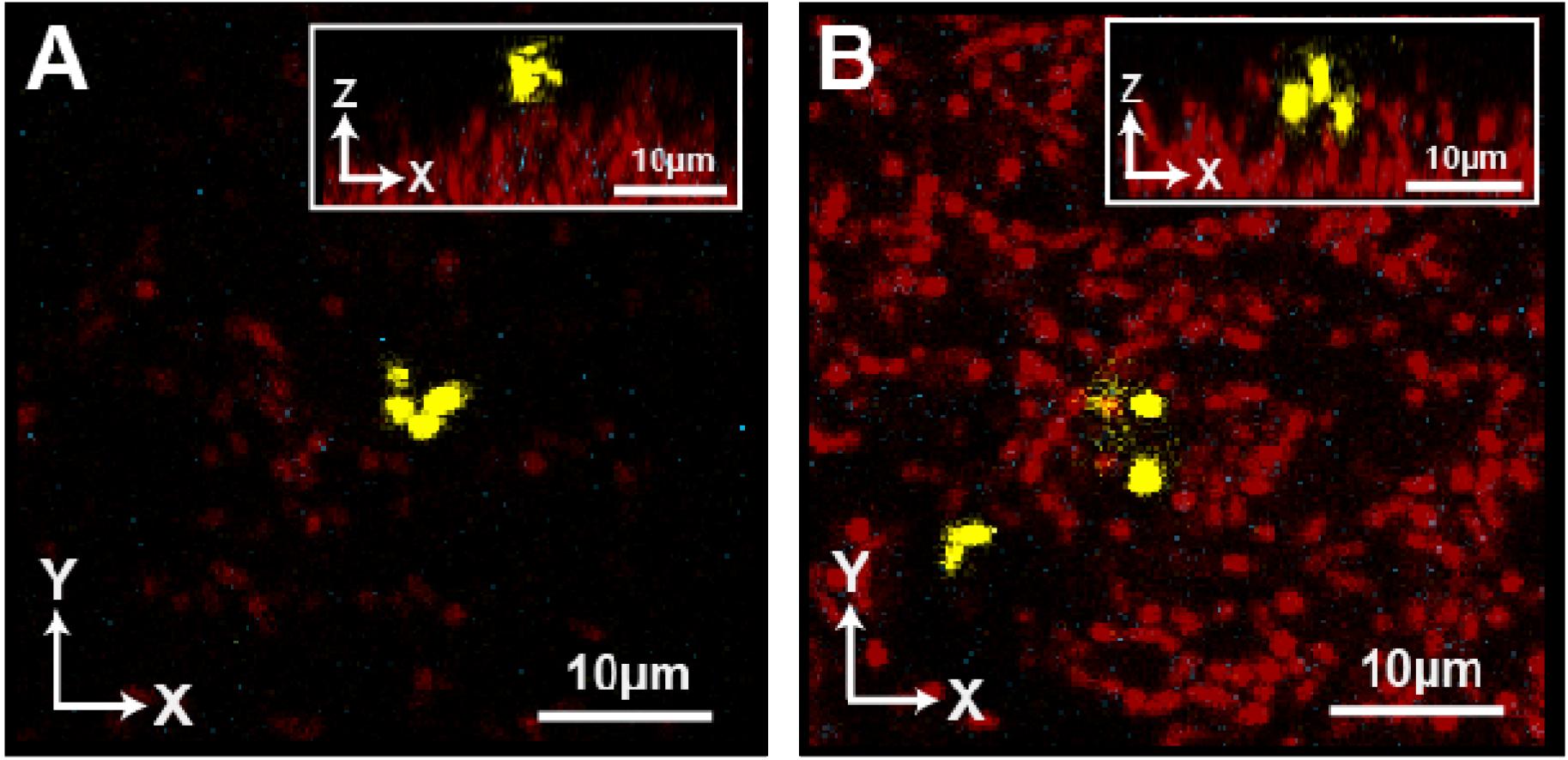
Invading Cells Do Not Produce Curli Matrix. (A) Representative image of invading cells (yellow) attached to resident biofilm (red) 3 hours following the conclusion of the invasion. Invading cells encode *csgA* with a C-terminal 6x-His tag. No matrix was detectable by immunostaining. (B) Representative image of invading cells ten hours after invasion. Again, invaders were not producing curli matrix.

**Figure S4.**
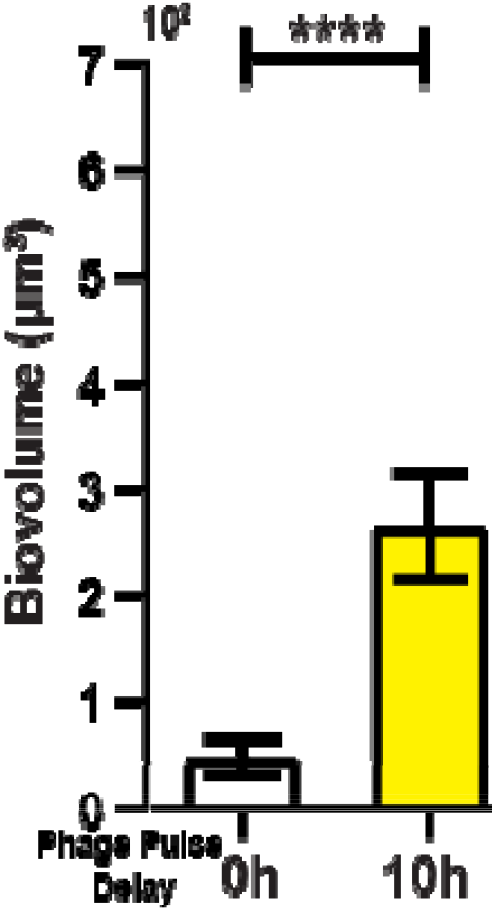
Visualization and quantification of invasion success of *E. coli* lacking the CsgB baseplate required for curli polymerization, with phage exposure post-invasion. (A) Average invading biovolume per field of view (150 μm x 150 μm x 15 μm; L, W, H). Error bars represent standard error of the mean. Pairwise comparisons were performed with Mann-Whitney Signed Ranks tests with Bonferroni correction (*n*=7; **** denotes *p*<0.00005). (B) Invading cells are killed when phages are introduced immediate after their arrival. (C) Invading cells are not killed when phages are introduced immediate after their arrival.

